# CSGDA: A Cell State-Guided Graph Domain Adaptation Network for Single-Cell Drug Response Prediction

**DOI:** 10.64898/2026.07.02.735966

**Authors:** Fen Yan, Xiyue Cao, Feiqiao Mao, Zhuhong You, Yitao Chen, Zhihua Du, Yu-An Huang

## Abstract

Intratumoral heterogeneity drives cancer recurrence and metastasis, yet single-cell drug response prediction faces severe “cross-domain” challenges, such as applying in vitro models to in vivo tissues or inferring metastatic resistance from primary tumors. These scenarios trigger distribution shifts arising from heterogeneous sequencing platforms, distinct tissue microenvironments, and metastatic evolution—problems rarely addressed by existing methods. We introduce CSGDA, a cell state-guided graph domain adaptation framework designed to predict drug responses across these biological heterogeneities. CSGDA incorporates biological priors to map gene expression into functional cell states, guiding a structure learning module to construct robust cell topology. To conquer distribution shifts, the model employs graph domain adaptation combined with a novel overlap penalty mechanism. Extensive benchmarks on five scRNA-seq datasets demonstrate that CSGDA outperforms state-of-the-art methods, achieving an average gain of ∼6% in ACC and AUPR. Beyond prediction accuracy, we employed integrated gradients to effectively pinpoint key genes involved in drug resistance within a challenging cross-metastasis cisplatin dataset. These findings underscore CSGDA’s superior performance in single-cell drug response prediction and its potential in resolving single-cell heterogeneity, paving the way for precision medicine.

## 1. Introduction

Intratumoral heterogeneity constitutes the molecular basis of drug resistance and recurrence (Marusyk *et al*., 2020; Ghorbian, 2025), yet the homogenized profiles derived from traditional Bulk RNA-seq inevitably mask critical rare subpopulations. Although scRNA-seq offers an unprecedented single-cell perspective for unraveling tumor microenvironment evolution and resistance mechanisms (Huang *et al*., 2023), reliance solely on wet-lab experiments to construct comprehensive drug response atlases lacks economic viability and scalability in clinical implementation due to prohibitive costs and complex protocols (Van de Sande *et al*., 2023).

Existing methods for predicting drug reactions, such as SCAD (Zheng *et al*., 2023), scDEAL (Chen *et al*., 2022), scAdaDrug (Liu *et al*., 2025), SSDA4Drug (Huang & Liu, 2025) and scGSDR (Huang *et al*., 2025), primarily adhere to a “Bulk-to-Single-Cell” cross-modal transfer learning paradigm, utilizing large-scale cell line population data as the source domain to transfer drug response knowledge to single-cell data (Maeser *et al*., 2024; Yan *et al*., 2025). However, this paradigm faces dual challenges in real-world scenarios. The first is Inherent Manifold Structure Mismatch. From a geometric perspective, Bulk data converges to the “centroid of a convex hull,” representing smoothed population averages, whereas scRNA-seq data constitutes highly non-linear “non-convex manifolds.” Forcing the alignment of such topological heterogeneity often results in subtle drug response features being overshadowed by statistically significant batch effects or broad cell-type main effects (Hui *et al*., 2024). The second is complex non-I.I.D. shifts. Multi-level distribution deviations introduced by discrepancies in sequencing platforms and microenvironments severely compromise the stability of feature alignment, leading to decision boundary drift (Park *et al*., 2021). Consequently, existing methodologies are largely confined to idealized or homologous contexts, failing to incorporate highly heterogeneous real-world clinical scenarios, such as cross-tissue adaptation and cross-metastasis evolution, which are essential for modeling the true complexities of actual patient data(Gundem *et al*., 2015; Savino *et al*., 2025).

With the rapid accumulation of drug-related scRNA-seq data, we advocate a shift towards an end-to-end “Single-cell to Single-cell” transfer paradigm and propose CSGDA (Cell State-Guided Graph Domain Adaptation). CSGDA is a general graph domain adaptation framework tailored for high-noise, non-I.I.D. data, with cell states serving as the core semantic variables. Instead of directly aligning high-dimensional noisy gene expression, CSGDA innovatively leverages “cell functional states” (e.g., EMT, DNA repair)—which possess intrinsic cross-domain invariance—as semantic anchors to construct a robust bridge connecting genotypes to phenotypes.

CSGDA achieves a deep integration of biological priors and graph structure learning through an end-to-end architecture. First, the model employs a biological prior-driven feature reconstruction mechanism that compresses the high-dimensional, sparse scRNA-seq gene space into low-dimensional ‘state manifolds’ with explicit biological significance, establishing interpretability at the input level. To mitigate the issue of spurious connections caused by high technical noise in scRNA-seq, we design a Cell State-Aware Graph Structure Learning (CSA-GSL) module. Unlike purely data-driven methods such as ProGNN (Jin *et al*., 2020) that may suffer from blind optimization in the absence of domain knowledge, CSA-GSL leverages cell state consistency as a prior constraint. By combining low-rank and sparsity restrictions, it dynamically reconstructs the relational topology between cells and utilizes a shared Graph Transformer to capture cross-domain consistent global patterns. Furthermore, to address the “ambiguous intermediate states” prevalent in non-I.I.D. scenarios, CSGDA incorporates an Overlap-aware Penalty strategy within its adversarial domain adaptation mechanism. By explicitly penalizing the overlapping regions of cross-domain category distributions, this strategy synergistically enlarges inter-class margins and stabilizes decision boundaries, thereby ensuring the precise identification of easily confused, drug-resistance-determining rare cell subpopulations.

CSGDA was extensively benchmarked on five diverse scRNA-seq datasets spanning varied platforms and metastatic stages. Experimental results demonstrate that CSGDA outperforms state-of-the-art methods in these highly heterogeneous transfer scenarios. Beyond the enhancement in predictive accuracy, a key advantage of CSGDA lies in its biological interpretability, enabling the precise identification of key biomarkers associated with drug-resistant phenotypes through unsupervised feature attribution analysis. Overall, CSGDA not only provides a solution for single-cell drug response prediction but also establishes a general paradigm for the cross-domain modeling of noisy single-cell data.

## 2. Materials and methods

### 2.1. Datasets

To comprehensively evaluate the robustness and generalizability of CSGDA across diverse biological heterogeneities, five representative single-cell RNA-sequencing (scRNA-seq) datasets were utilized in this study. These datasets encompass a variety of challenging cross-domain scenarios, including cross-platform, cross-cell line, cross-tissue (monotherapy and combination therapy), and cross-metastasis adaptation.

### 2.2. Cell State-Guided Graph Structure Learning Module

The inherent high-dimensional sparsity and dropout noise characteristic of scRNA-seq data frequently result in cell relationship graphs compromised by spurious connections or lacking critical biological associations. To reconstruct a faithful intercellular topology, we introduce the CSA-GSL module (Fig. 1d). This module integrates biological prior knowledge to construct an initial Original Noisy Graph and utilizes a ProGNN-based optimization framework to denoise it.

**Figure 1.**
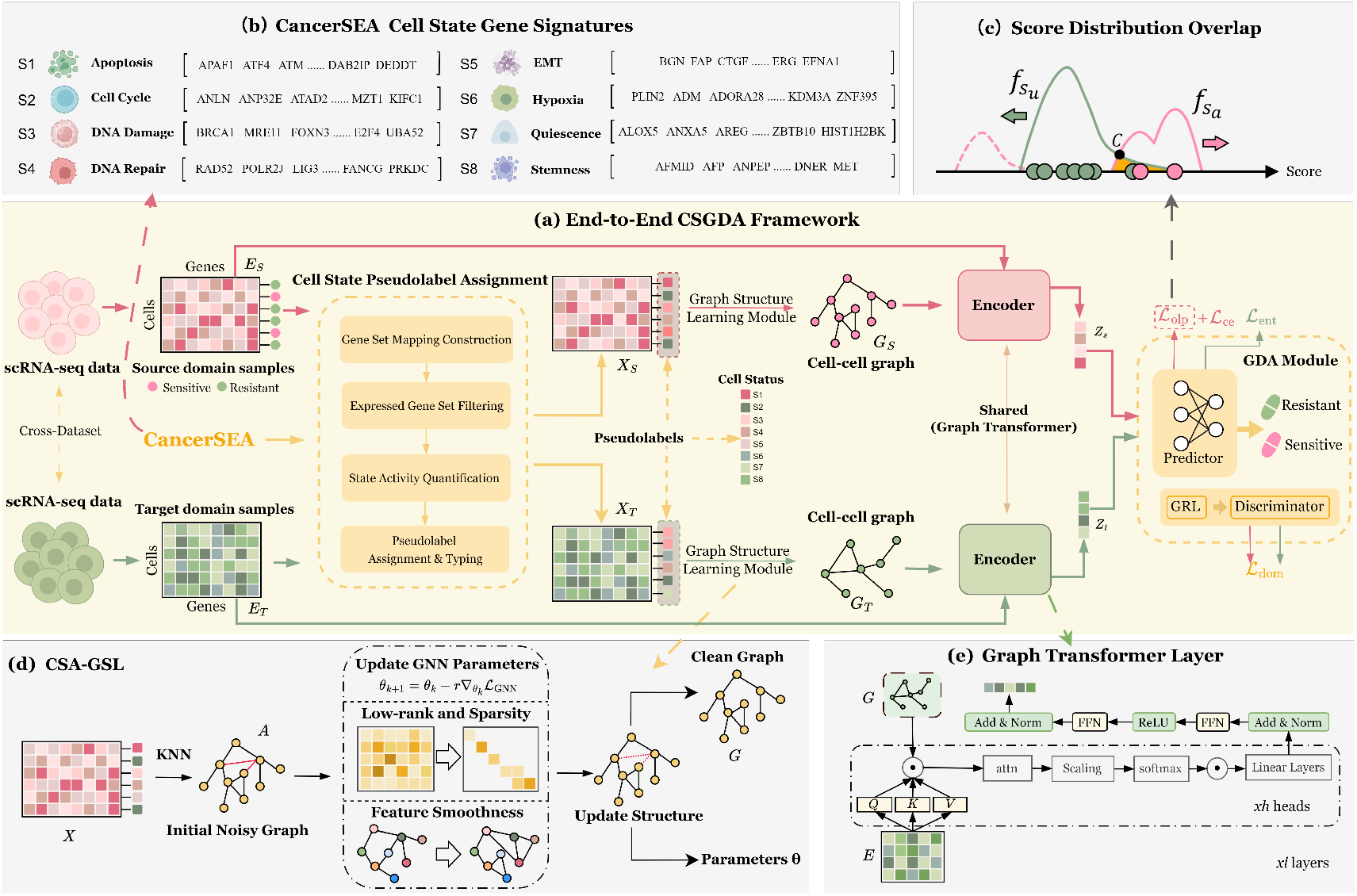
The architecture of CSGDA. The framework comprises (a) the complete End-to-End CSGDA Framework for cross-domain drug response prediction, (b) biological prior construction, (c) the GDA Module incorporating an overlap penalty mechanism, (d) the CSA-GSL Module for robust topology, and (e) the Shared Graph Transformer Encoder

#### (1) Cell State Extraction and Pseudo-Labeling

As depicted in Fig. 1b, the assignment proceeds through distinct steps:

*Construction of state-gene mapping*. Leveraging the CancerSEA database, we established a state-gene mapping dictionary 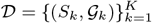, selecting *K* = 8 core functional states highly correlated with drug resistance.

*Expression gene filtering and cell state activity quantification*. We transform the preprocessed gene expression matrix *E ∈*ℝ^*N×M*^ into a biologically interpretable feature space,where *N* and *M* denote the number of cells and genes, respectively. Specifically, for each cell *i* and functional state *k*, we compute the state activity score by averaging the expression levels of genes shared between the state-specific marker set *G*_*k*_ and the detected gene set _data_. This process generates the dense cell state feature matrix *X ∈* ℝ^*N×*8^, effectively mitigating dropout effects.

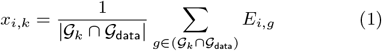

Here, *x*_*i,k*_ denotes the state activity score of cell *i* for the *k*-th state, where *E*_*i,g*_ represents the expression level of gene *g* in cell *i*. _*k*_ and _data_ correspond to the marker gene set for the *k*-th state and the total gene set in the scRNA-seq dataset, respectively, with _*k* data_ indicating the count of valid overlapping marker genes.

*Pseudo-label assignment*. To facilitate optimization, we derive pseudo-labels from the computed state activity scores. Adopting the “Winner-Takes-All” principle, we assign the dominant state pseudo-label 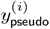 to the state exhibiting the maximal score. These labels serve as biological guidance for interpreting cellular heterogeneity.

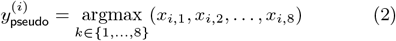

Here, 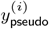 denotes the assigned dominant state pseudo-label for cell *i. x*_*i,k*_ denotes the state activity score for cell *i* under the *k*-th functional state. The search space *{*1, …, 8*}* corresponds to the set of eight core functional cell states defined in our framework.

#### (2) Construction of the Original Noisy Graph

Leveraging the feature matrix *X*, we construct the initial cell affinity graph *G*_init_ = (*A, X*) via the KNN algorithm. Although derived from semantic features, _init_ is designated as “noisy” as it retains spurious connections from technical noise.We define the adjacency matrix *A ∈ {*0, 1*}*^*N ×N*^ based on the *k*-nearest neighbor set *N*_*k*_(*v*_*i*_), where each entry *A*_*ij*_ is formulated as:

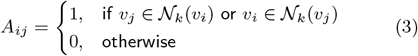

#### (3) Clean Graph Structure Learning

We employ a joint optimization strategy rooted in the ProGNN (Jin *et al*., 2020) framework, the workflow of which is shown in Fig. 1d, to learn an optimal graph structure *G* and GNN parameters *θ*. The overall loss function *L*_GSL_(*G, θ*) integrates three components: Classification guidance loss (*L*_GNN_) minimizes prediction error; Graph regularization loss (*L*_Reg_) imposes sparsity, low-rank, and feature smoothness constraints to enforce robustness; and a Fidelity constraint (*L*_Fid_) anchors the learned graph to the initial topology to prevent structural collapse.

We linearly combine the aforementioned terms to define the overall loss function *L*_GSL_(*G, θ*):

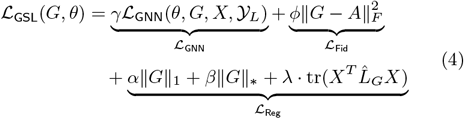

Graph refinement incorporates a sparsity constraint *α∥G∥*_1_ (*α* = 5 *×* 10^*−*4^) and a low-rank constraint *β∥G∥*_*∗*_ (*β* = 1.5). The hyperparameter *ϕ* balances reconstruction freedom against prior topology to preserve critical neighborhoods during denoising. Crucially, we introduce a feature smoothness constraint 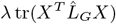(*λ* = 0.1) based on the homophily assumption of biological networks, where cells with similar biological states tend to connect.

By employing the normalized graph Laplacian 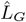 this constraint penalizes edges between cells that exhibit significant differences in their 8-dimensional state features *X*. This mechanism effectively filters out spurious cross-subpopulation associations while mitigating the impact of varying cellular connectivities on feature smoothness. The normalized graph Laplacian 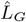 is defined based on the degree matrix *D ∈* R^*N×N*^ of the clean graph *G*. Here, *D* is a diagonal matrix where each diagonal element represents the number of adjacent cells, corresponding to the cellular connectivity within the single-cell microenvironment:

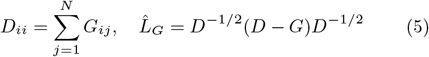

To ensure that the normalized feature differences between connected cells are minimized, the trace form of the feature smoothness loss can be transformed into an intuitive summation

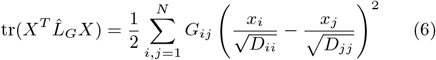

Finally, the hyperparameter *γ* weights the classification loss against these structural constraints. To solve this overall objective, we employ an alternating optimization strategy to iteratively update both the graph structure *G* and the GNN parameters *θ*.

#### a. Graph structure update

Under the premise of fixing the current GNN parameters *θ*, we update the graph structure *G* by minimizing *L*_GSL_. This step enforces the graph regularization constraints on *G*. Given that the sparsity (*∥G∥* _1_) and low-rank (*∥G∥*_*∗*_) terms are non-smooth, we adopt the Forward-Backward Splitting method. Specifically, we sequentially handle the non-smooth terms of the nuclear norm and the *L*_1_ norm using incremental proximal descent. First, we compute the gradient for all differentiable terms and perform gradient descent. Then, we sequentially apply the proximal operators of the nuclear norm and the *L*_1_ norm to enforce the low-rank and sparsity constraints. Finally, a projection operator is applied to ensure the validity of the graph. The specific process is as follows:

#### Gradient descent update

We compute the gradient of the differentiable terms related to *G* in the total loss and perform a gradient descent update:

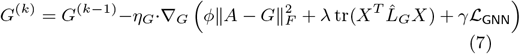

Here, *k* is the iteration round, *η*_*G*_ = 0.01 is the dedicated learning rate for updating the graph structure *G*, and _*G*_ is the gradient operator with respect to *G*.

#### Low-rank structure recovery

We apply the nuclear norm proximal operator to the gradient-updated *G*^(*k*)^ to enforce the low-rank constraint:

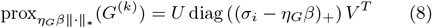

Here, *G*^(*k*)^ = *U* diag(*σ*_*i*_)*V* ^*T*^ is the singular value decomposition (SVD) of *G*^(*k*)^, *σ*_*i*_ is the *i*-th singular value of *G*^(*k*)^, and (*x*)_+_ = max(*x*, 0) is the positive part function. The singular values are reduced through soft-thresholding to achieve the low-rank property of the graph.

#### Sparsification filtering

We apply the *L*_1_ norm proximal operator to the result of the low-rank constraint to enforce the sparsity constraint:

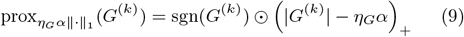

Here, sgn(*G*^(*k*)^)_*ij*_ is the sign matrix (taking 1 when 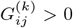, 0 when 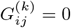, and *−*1 when 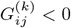), and *⊙* denotes t Hadamard product (element-wise product). The spurious edges with low weights are set to 0 through soft-thresholding.

#### Projection operator constraint

After each update, a projection operator *P*_*S*_(*G*) is used to ensure the symmetry and weight validity of the graph:

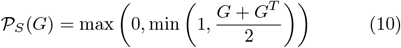

Here, 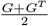 performs the symmetry projection to ensure the graph is undirected, and max(0, min(1, *·*)) performs the numerical range projection to preserve the physical meaning of the weights.

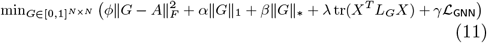

#### b. GNN parameter update

Fixing the updated graph structure *G*, we optimize the GNN parameters *θ* via standard gradient descent to minimize the classification loss:

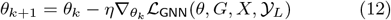

where *Y*_*L*_ denotes the ground-truth drug response labels for source domain samples, and *η* represents the learning rate. This procedure guarantees that the learned graph *G* delivers the most effective topology for downstream node classification.

By alternating between these two steps until convergence, we obtain a robust, denoised graph structure *G* that significantly enhances drug response prediction. This refined topology serves as the structural input for the subsequent shared Graph Transformer encoder.

### 2.3. Shared Graph Transformer Encoder

To overcome distribution shifts arising from varying platforms, tissue of origin, or disease states, we engineered a weight-shared Graph Transformer encoder, *E*_sh_, the architecture of which is detailed in Fig. 1e. This module is designed to extract domain-invariant feature representations. Prior to the Transformer layers, we map high-dimensional sparse gene expression features into a continuous latent space via a fully connected layer to enhance feature expressiveness. For cell *i*, the initial feature embedding 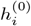 is computed as:

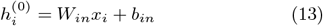

where *x*_*i*_ *∈*ℝ^*M*^ denotes the raw gene expression vector, *W*_in_ *∈* ℝ^*d×M*^ is the projection matrix, and *b*_in_ ∈ R^*d*^ is the bias vector.

The shared encoder processes inputs from both domains—comprising the gene expression matrix *E* and the homogeneous graph structure *G* derived from the CSA-GSL module—through an identical architecture. The forward propagation is formalized as:

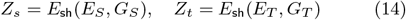

where *E*_*S*_, *E*_*T*_ represent input expression matrices, and *Z*_*s*_, *Z*_*t*_ *∈* ℝ^*N×d*^*h* denote the resulting domain-invariant embeddings.*E*_sh_ comprises stacked Graph Transformer layers, the detailed architecture of which is presented in Fig. 1e.

To fully exploit biological neighborhoods in *G*, we designed a structure-constrained multi-head attention mechanism. Unlike standard global attention, computation is strictly confined to local neighborhoods defined by *G*. For any cell *i*, the feature update at layer *l* + 1 proceeds as follows:

#### Structure-Constrained Attention

We calculate Query, Key, and Value vectors and filter noisy connections using *G*:

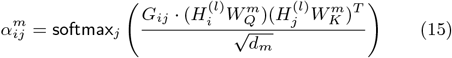

Here, 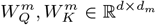 are projection matrices. The scaling factor 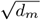 prevents gradient vanishing caused by large dot products. *G*_*ij*_ acts as a structural mask: non-zero entries enable dynamic weighting based on expression similarity, while zero entries physically suppress connections, ensuring aggregation is exclusive to homogeneous neighbors.

#### Feature Aggregation

To mitigate training instability caused by scRNA-seq sparsity, we employ Batch Normalization (BN) after the residual connection:

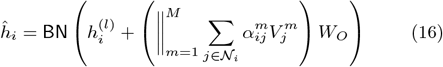

where 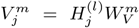 is the value vector, 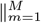 denotes concatenation across *M* heads, and 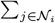 performs weighted aggregation within the neighborhood *N*_*i*_. *W*_*O*_ represents the output linear projection matrix, which maps the concatenated multi-head features back to the target dimension.

#### Feed-Forward Update

The intermediate features undergo non-linear transformation via a Feed-Forward Network (FFN), followed by a second residual and BN step:

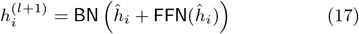

This “aggregation-update” mechanism enables the encoder to extract robust representations that combine local topology with global semantics. To prevent feature degradation in deep networks, a global residual connection is applied, adding the initial projected features to the output of the final Graph Transformer layer.

#### Stochastic Latent Regularization

To mitigate the high-dimensional sparse noise in scRNA-seq data, we incorporate a stochastic latent embedding strategy. Instead of deterministic mapping, we employ parallel layers to estimate the mean and log-variance log(*σ*^2^) of the cellular latent distribution. The final embedding Z isgenerated via the reparameterization trick:

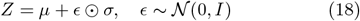

### 2.4 Graph-based Domain Adaptation Module

To transfer predictive efficacy from the source domain to an unlabeled target domain, we developed a GDA module rooted in adversarial learning(Fig. 1a). Furthermore, to address class overlap, we employ a specific penalty mechanism (Fig. 1c). This module mitigates distribution shifts arising from sequencing platform variability or tissue heterogeneity.

#### Drug Response Predictor

Given cell embedding *z*_*i*_, the predictor generates the class probability distribution *ŷ*_*i*_ (sensitive vs. resistant) using a linear projection and Softmax function.

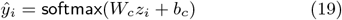

#### Adversarial Domain Alignment

To address covariate shift, we employ an adversarial strategy where a domain discriminator *D* distinguishes source from target embeddings, while the shared encoder *E*_sh_ generates domain-invariant features. The adversarial loss *L*_dom_ is a binary cross-entropy objective:

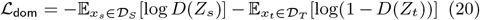

Here, *D*_*S*_ and *D*_*T*_ denote source and target distributions, and *Z*_*s*_ and *Z*_*t*_ are embeddings from *E*_sh_. The discriminator *D*(*·*) outputs the probability of belonging to the source domain (label 1) versus the target (label 0), with E denoting expectation. To implement the minimax game, we introduce a Gradient Reversal Layer (GRL), defined as a pseudo-function *R*(*Z*) that performs identity mapping (*R*(*Z*) = *Z*) during forward propagation but reverses gradients during backpropagation. This enforces the following update rule for encoder parameters *θ*_enc_:

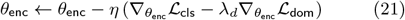

Here, *θ*_enc_ and *η* denote the encoder parameters and learning rate, respectively. 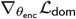 represents the gradient of the domain loss. The coefficient *−λ*_*d*_ induces gradient ascent to maximize the discriminator’s error, where *λ*_*d*_ controls the weight of this adversarial alignment to eliminate non-biological batch effects.

#### Target Domain Information Entropy Minimization

To enforce the cluster assumption, we minimize the entropy of target probabilities *P*_*t*_ = softmax(*C*(*Z*_*t*_)):

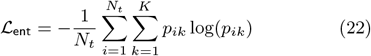

Here, *N*_*t*_ denotes the target cell count, and *p*_*ik*_ represents the probability of cell *i* belonging to class *k* (where *K* = 2). This constraint drives decision boundaries away from high-density data regions.

#### Classification Optimization Based on Overlap Penalty

While the standard cross-entropy loss guides source domain training, it often fails to guarantee sufficient inter-class margins. To enforce robust decision boundaries, we introduce an overlap loss (*L*_olp_). We first partition the prediction probabilities for the “Sensitive” class into two sets based on ground truth labels:

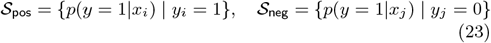

where *S*_pos_ contains scores for true sensitive cells and *S*_neg_ contains scores for true resistant cells. The overlap loss minimizes the intersection of these distributions, as illustrated in Fig. 1c, where the mechanism pushes the score distributions of sensitive and resistant cells apart:

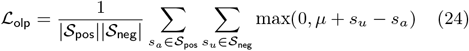

Here, *μ* is the preset margin threshold. Minimizing this loss imposes a dual constraint: suppressing resistant scores (*s*_*u*_ *∈ S*_neg_ *→* 0) while boosting sensitive confidence (*s*_*a*_ *∈ S*_pos_ *→* 1). The final source domain objective combines cross-entropy with this weighted overlap loss:

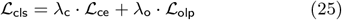

Here, *λ*_c_ and *λ*_o_ are fixed weights balancing source classification. This strategy delineates a wide-margin decision boundary, effectively tolerating feature perturbations induced by domain shifts.

### 2.5. Total Objective Function

To achieve robust cross-domain generalization, we optimize a unified framework defined by:

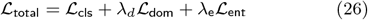

Specifically, the total objective function is composed of the source domain classification loss *L*_cls_ (integrating overlap penalty), the adversarial domain adaptation loss *L*_dom_, and the target domain information entropy loss *L*_ent_. To prevent early overconfidence, *λ*_e_ follows a linear growth schedule *λ*_e_(*t*) = 0.1 *t/T*_max_. For stability, the adversarial weight *λ*_*d*_ employs a dynamic warm-up strategy 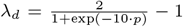 (where *p* is training progress); this initially suppresses domain alignment to prioritize biological feature learning, then gradually ramps up to enforce invariance.Upon convergence, auxiliary modules are discarded. The model operates efficiently using only the shared encoder and predictor: *ŷ* = softmax(*C*(*E*_sh_(*x, G*))).

## 3. Experiments

To distinguish the performance gains of our paradigm shift from those of CSGDA’s specific architecture, we first conducted a baseline comparison using CSGDA and six representative baseline models. As shown in Fig. 3 and Table S5, shifting from Bulk-to-SC to the SC-to-SC paradigm significantly circumvents manifold mismatches, providing a superior foundation. Building upon this validated paradigm, we next evaluate the cumulative improvements achieved by CSGDA’s architectural optimizations.

### 3.1. Analysis of Model Performance

To address the inherent class imbalance in scRNA-seq data, we evaluated CSGDA using AUROC, AUPR, ACC, and F1-macro, systematically comparing it against five state-of-the-art methods across five scenarios (Table 1). To ensure a fair and rigorous comparison, the hyperparameters for both our proposed CSGDA and all baseline methods were meticulously tuned. For comparison, we adopted the official implementations of the benchmark methods and followed their recommended or default hyperparameter settings. All methods were applied to the same preprocessed datasets under identical experimental conditions to ensure a fair and reproducible comparison. CSGDA achieved robust performance overall, particularly in the cross-platform scenario, where it attained near-perfect scores alongside scAdaDrug and scGSDR; UMAP visualization (Fig. 2A-B, left panel) attributes this to the high intrinsic separability of resistance and sensitivity classes in low-noise settings.

**Table 1.** Performance comparison between CSGDA and baseline models on five datasets.

| Method | Exp1: Cross-Platform |  |  |  | Exp2: Cross-Cell Line |  |  |  | Exp3: Cross-Tissue (Monotherapy) |  |  |  | Exp4: Cross-Tissue (Combination) |  |  |  | Exp5: Cross-Metastasis |  |  |  |
| --- | --- | --- | --- | --- | --- | --- | --- | --- | --- | --- | --- | --- | --- | --- | --- | --- | --- | --- | --- | --- |
|  | ACC | F1-macro | AUC | AUPR | ACC | F1-macro | AUC | AUPR | ACC | F1-macro | AUC | AUPR | ACC | F1-macro | AUC | AUPR | ACC | F1-macro | AUC | AUPR |
| scDEAL | 0.931 | 0.928 | 0.980 | 0.984 | 0.524 | 0.368 | 0.678 | 0.697 | 0.585 | 0.642 | 0.681 | 0.790 | 0.688 | 0.552 | 0.818 | 0.596 | 0.693 | 0.583 | 0.692 | 0.551 |
| SCAD | 0.992 | 0.992 | 0.998 | 0.998 | 0.871 | 0.890 | 0.882 | 0.914 | 0.831 | 0.811 | 0.911 | 0.859 | 0.834 | 0.817 | 0.899 | 0.824 | 0.794 | 0.798 | 0.849 | 0.885 |
| scADADrug | 0.958 | 0.962 | 1.000 | 1.000 | 0.541 | 0.666 | 0.696 | 0.850 | 0.839 | 0.785 | 0.906 | 0.857 | 0.863 | 0.846 | 0.898 | 0.767 | 0.554 | 0.713 | 0.399 | 0.497 |
| scGSDR | 0.984 | 0.984 | 0.999 | 0.999 | 0.673 | 0.672 | 0.793 | 0.834 | 0.674 | 0.611 | 0.720 | 0.597 | 0.576 | 0.473 | 0.533 | 0.419 | 0.564 | 0.412 | 0.615 | 0.701 |
| SSDA4Drug | 0.930 | 0.929 | 0.998 | 0.998 | 0.831 | 0.826 | 0.894 | 0.873 | 0.851 | 0.843 | 0.913 | 0.847 | 0.814 | 0.802 | 0.912 | 0.842 | 0.782 | 0.774 | 0.850 | 0.899 |
| CSGDA | 0.994 | 0.994 | 0.999 | 0.999 | 0.891 | 0.890 | 0.931 | 0.952 | 0.857 | 0.846 | 0.924 | 0.885 | 0.912 | 0.907 | 0.970 | 0.964 | 0.882 | 0.873 | 0.966 | 0.957 |

**Figure 2.**
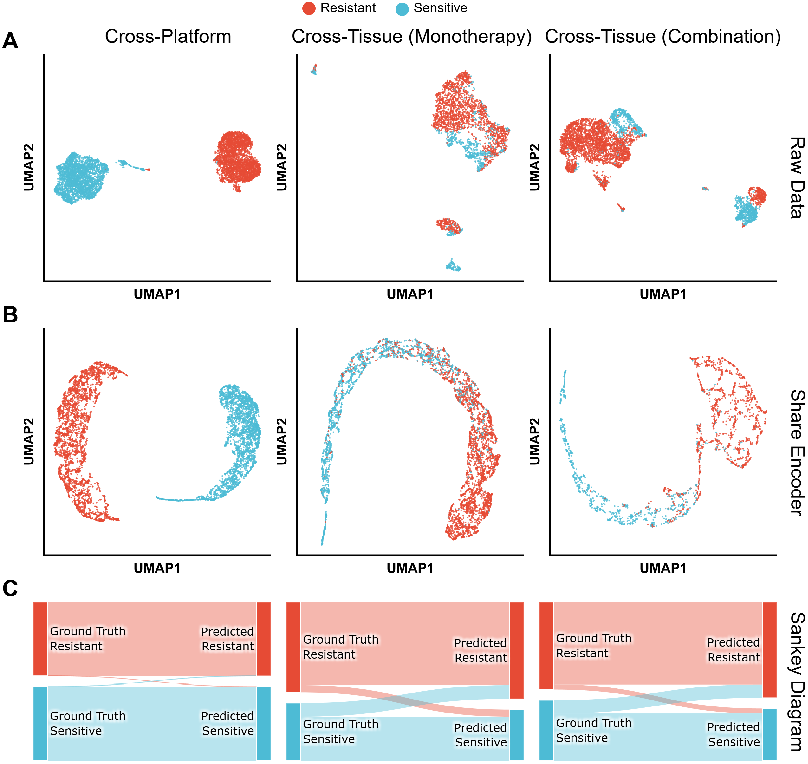
Feature alignment and prediction visualization. **(A-B)** UMAP comparison of **Raw Data (A)** and **Shared Encoder (B)** features across Cross-Platform, Monotherapy, and Combination tasks. **(C)** Sankey diagrams showing prediction consistency.

**Figure 3.**
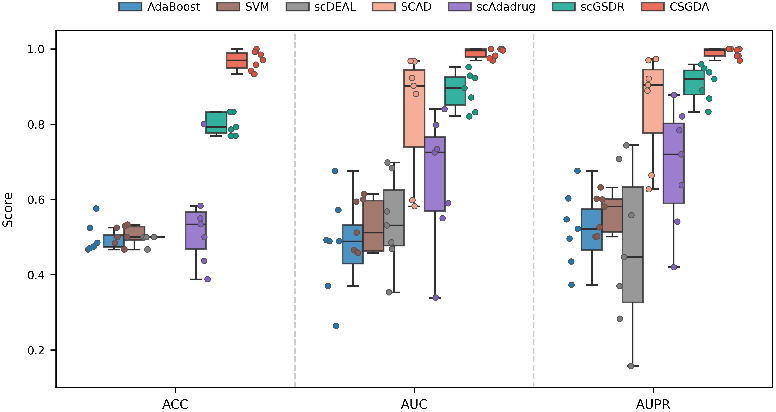
Overall performance distribution of CSGDA and six baseline models under the traditional Bulk-to-SC transfer paradigm.

However, as the scenarios progressed towards higher heterogeneity, performance stratification emerged, revealing the limitations of baseline architectures. First, unlike scDEAL which relies on statistical alignment (e.g., MMD), CSGDA employs a **Cell State-Guided** strategy. By aligning domains within a biological functional space rather than a purely statistical one, CSGDA better maintains performance when facing complex biological shifts. Second, while SCAD and scAdaDrug process cells as independent instances, CSGDA utilizes **Dynamic Structure Learning**. This allows it to capture non-linear topological relationships and intercellular dependencies, which are critical for identifying resistance in highly heterogeneous environments. Among the baselines, SSDA4Drug exhibited the strongest competitiveness. It is important to note that SSDA4Drug is a semi-supervised method that relies on partial target domain labels to guide adaptation. In contrast, CSGDA operates under a strictly Unsupervised Domain Adaptation (UDA) paradigm, with no access to target labels. Despite this “information disadvantage,” CSGDA surpassed SSDA4Drug in the complex scenarios.

UMAP visualization reveals the discriminative power of CSGDA’s latent embeddings (compare Fig. 2A and B). In the Cross-Platform task, CSGDA effectively preserved clear class boundaries and compressed intra-class distances (Fig. 2B, left panel), confirming its robustness in baseline tasks. Crucially, in the high-noise Cross-Tissue(Monotherapy) scenario, the model transformed entangled raw data into a distinct “Inverted U-shape Manifold” (Fig. 2A-B, middle panel). This topology implies a continuous biological gradient rather than discrete clustering, suggesting that CSGDA captures the phenotypic plasticity and intermediate states traversing from sensitivity to resistance. Similarly, in the complex Cross-Tissue(Combination) setting, the model resolved irregular distributions into a compact “Hook-like Topology” (Fig. 2B, right panel), demonstrating strong non-linear feature extraction guided by cell states. Sankey diagrams (Fig. 2C) further quantify this alignment, showing high consistency between predictions and ground truth, particularly for the challenging samples in these cross-tissue scenarios.

### 3.2. Ablation Study

To validate the contribution of each component in CSGDA, we conducted systematic ablation studies across five experimental scenarios. We compared the full model against variants lacking biological priors, dynamic graph learning, domain adaptation, and hybrid loss functions, respectively. As illustrated in the comprehensive performance distributions (Fig. 4), CSGDA demonstrated superior performance across most experimental scenarios and metrics, validating the synergy of the proposed modules.

**Figure 4.**
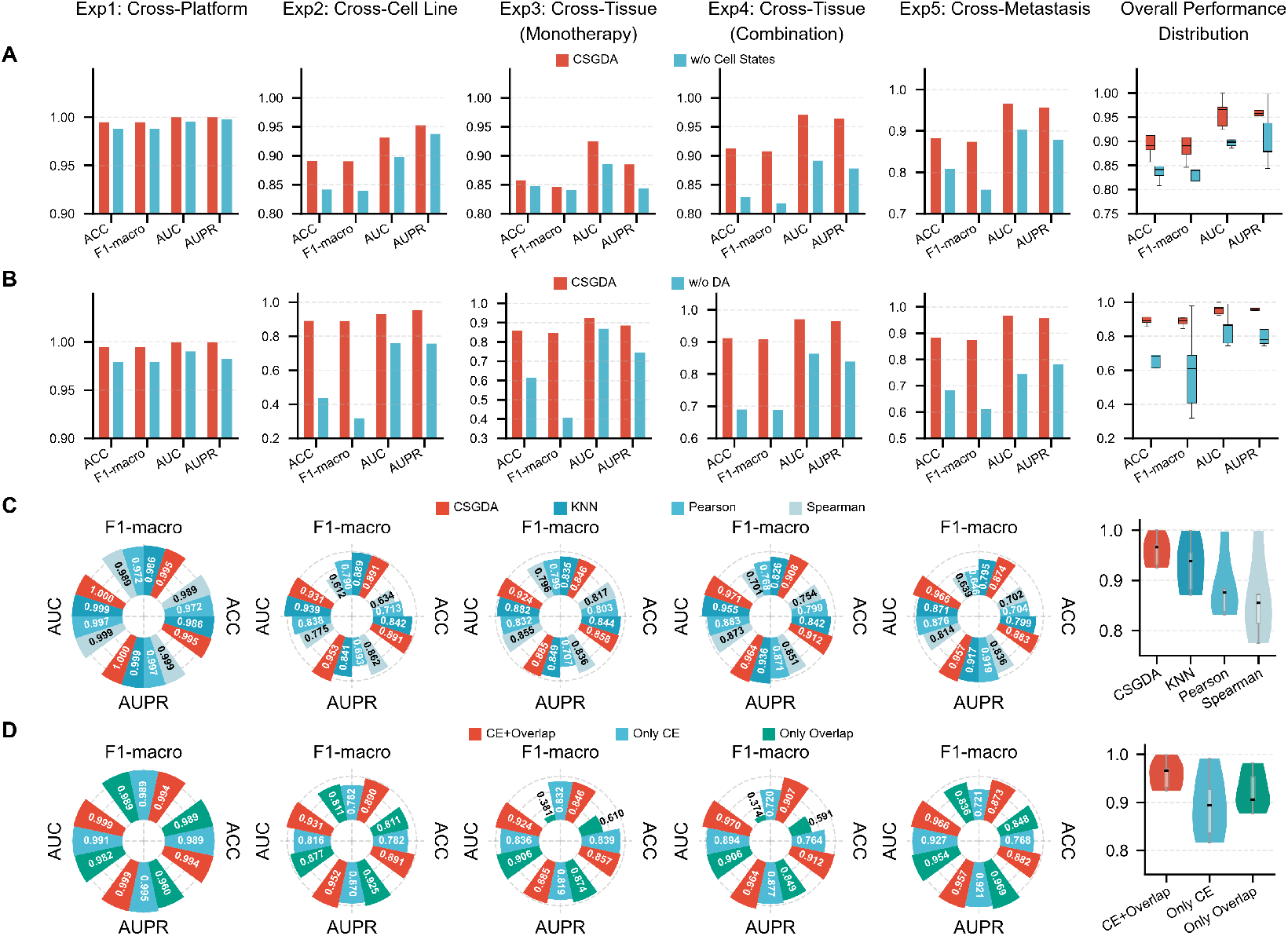
Comprehensive ablation study of CSGDA components across five experimental scenarios. (A) Impact of biological priors. (B) Contribution of the domain adaptation module. (C) Performance comparison of the CSA-GSL module against static graph construction strategies. (D) Efficacy of the hybrid loss function strategy.

#### Impact of Biological Priors

We first verified the superiority of introducing cell states over purely data-driven features. Compared to the variant using PCA and K-Means (w/o Cell States), CSGDA exhibits significant robustness in complex environments(Fig. 4A). For instance, in the highly heterogeneous cross-tissue scenario, CSGDA maintained a high AUC of *∼*0.97, whereas the statistical feature variant dropped to 0.89. This disparity confirms that biological priors provide conservative feature anchors (e.g., DNA repair semantics), effectively suppressing the model from learning spurious correlations from stochastic scRNA-seq noise.

#### Effectiveness of CSA-GSL

We then assessed the graph learning module against static strategies (KNN, Pearson, Spearman), as illustrated in (Fig. 4C). Static methods failed to capture complex dependencies in the cross-metastasis scenario, with AUC scores stagnating below 0.88. In contrast, CSA-GSL achieved a remarkable AUC of 0.966. This improvement stems from the shift from “static feature similarity” to “dynamic semantic consistency,” where the graph structure is iteratively reconfigured based on updating predictions, correcting erroneous connections caused by dropout events.

#### Contribution of Domain Adaptation

The exclusion of the domain adversarial network (w/o DA) precipitated a drastic performance collapse in cross-domain tasks(Fig. 4B). Specifically, in the combination therapy scenario, the absence of DA caused the AUC and AUPR to regress sharply from 0.97 and 0.964 to 0.863 and 0.839, respectively. This gap (*>* 0.1) demonstrates that domain alignment serves as a crucial bridge, mitigating inter-domain distribution discrepancies (e.g., tissue origins) to ensure reliability on unseen targets.

#### Necessity of Cross-Entropy and Overlap Loss

Finally, Fig. 4D highlights the limitations of single-modal losses. Relying solely on overlap loss (*L*_olp_) in Exp. 4 resulted in a random-level F1-macro (0.374) despite high ranking metrics, due to the lack of probability calibration. Conversely, using only cross-entropy (*L*_olp_) in the tumor evolution scenario (Exp. 5) led to overfitting source boundaries. By organically integrating absolute probability supervision (*L*_olp_) with relative topological constraints (*L*_olp_), CSGDA achieves peak F1-macro scores (e.g., 0.873 in Exp. 5), ensuring both decision boundary precision and generalization.

### 3.3. Biological Interpretability Analysis

To assess whether the CSGDA model captures genuine biological mechanisms underlying drug resistance, we employed the Integrated Gradients (IG) algorithm to establish a multi-level “Drug Response–Gene–State” validation framework. Here, we focused on the challenging cross-metastasis scenario predicting Cisplatin response (Experiment 5) as our representative case study. Our core logic involved identifying the genes that contribute most to resistance or sensitivity predictions (based on IG Scores), examining the enrichment patterns of these key genes within biological cell states, and subsequently inferring the biological mechanisms driving the model’s decisions.

#### (1) Identification and Validation of Driver Genes

We first analyzed the core gene expression patterns that drive the model to predict resistance. As shown in Supplementary Table S6, the top 20 high-contribution genes identified by the model exhibited substantial IG scores, suggesting that the model mathematically regards the high expression of these specific genes as a significant determinant of cellular drug resistance. This computational finding aligns closely with clinical observations. Notably, RAD51C was identified as the top-ranking driver. As a paralog of RAD51, RAD51C is an essential component of the homologous recombination repair pathway; it assists in the assembly of the RAD51 recombinase on single-stranded DNA and plays a crucial role in repairing cisplatin-induced double-strand breaks (Hoppe *et al*., 2021). Clinical studies have indicated that elevated RAD51C expression is closely associated with cisplatin resistance and poor prognosis in various cancers, including ovarian and oral squamous cell carcinomas (Jiang *et al*., 2024; Galluzzi *et al*., 2012). The model’s assignment of high weight to RAD51C suggests it is detecting genuine resistance drivers rather than relying on data noise.

#### (2) Cell State Mapping and Resistance Mechanisms

We utilized the CancerSEA database as a source of biological prior knowledge to bridge the gap between “micro-level genes” and “macro-level cell states.” Specifically, we employed gene signature sets from eight cancer-related functional states (e.g., DNA Repair, Hypoxia) as a mapping dictionary. By matching the high-contribution genes identified by CSGDA with this dictionary and calculating the IG scores for genes within each state, we quantified the overall contribution of different cell states to the resistance prediction (Fig. 5). This mapping analysis revealed a distinct pattern: a majority of high-contribution genes (such as RAD51C and ERCC1) aligned with the DNA Repair and DNA Damage states. Consequently, at the macro-state level, the DNA repair mechanism exhibited a predominant contribution. This result is consistent with the pharmacological mechanism of cisplatin, which aims to eliminate cancer cells via DNA damage, prompting resistant cells to upregulate DNA repair pathways to survive.

**Figure 5.**
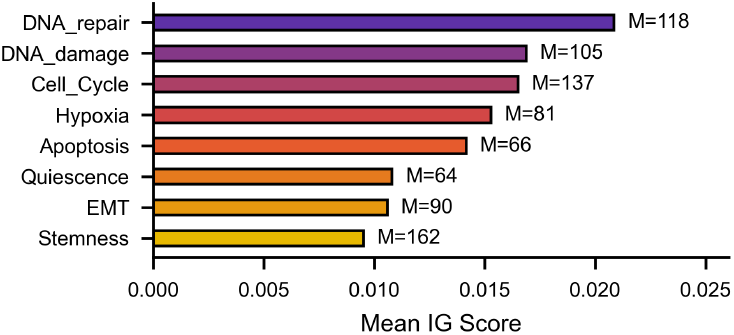
Global importance ranking of functional states. The bar chart displays the mean Integrated Gradients (IG) scores of high-contribution genes across eight predefined CancerSEA states.

Additionally, the model identified Cell Cycle and Hypoxia as secondary factors. Cisplatin-induced damage typically triggers G2/M phase arrest, providing a window for DNA repair.

Resistant cells often possess flexible cell cycle regulation, allowing them to evade apoptosis signals and rapidly resume proliferation (Olszewska *et al*., 2022; Sarin *et al*., 2017). Hypoxia within tumors can induce HIF-1*α* expression, which reshapes metabolic modes (e.g., by enhancing glycolysis) and transmits resistance signals to surrounding cells via mechanisms such as exosomes (Wang *et al*., 2021). The model’s detection of hypoxia reflects an understanding of complex resistance patterns driven by tumor microenvironmental stress. This indicates that CSGDA captures not only the primary drivers but also the auxiliary roles of tumor proliferation and microenvironmental pressure in drug resistance.

#### (3) Unsupervised Discovery Beyond Priors

A key advantage of this framework is its ability to discover features beyond the boundaries of predefined knowledge. As shown in Fig. 6, the model identified a group of significant high-contribution genes—including NDUFA11, NDUFS6, and TMEM205—in an unsupervised manner. Although these genes were not included in the initial CancerSEA dictionary, the model detected their strong contribution to resistance based on intrinsic data distributions.

**Figure 6.**
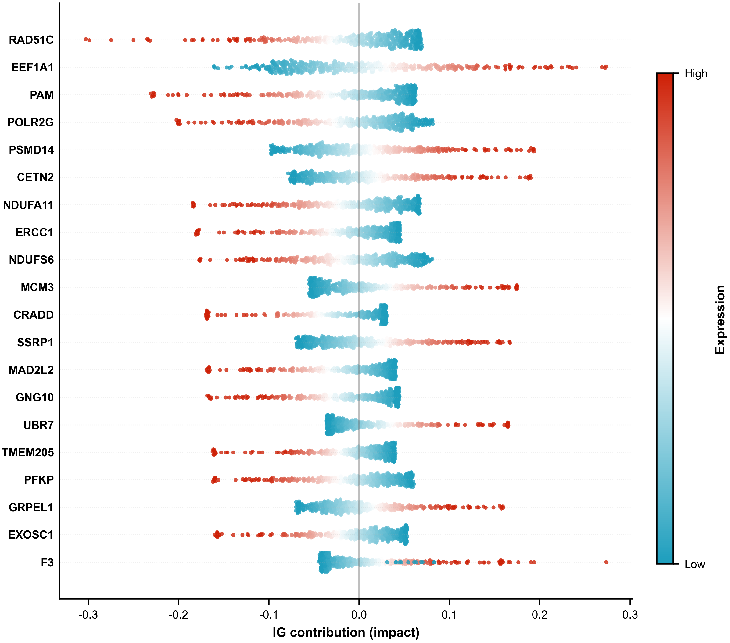
Beeswarm plot of top driver genes. The clustering of high-expression dots on the positive side demonstrates their strong driving force toward Cisplatin resistance.

The identification of TMEM205, in particular, offers strong biological support for the model’s predictive mechanism.

Literature confirms that TMEM205 is a 21 kDa transmembrane protein, primarily localized to the cell surface and the trans-Golgi network, and is specifically highly expressed in cisplatin-resistant cells (Shen *et al*., 2010). Its core resistance mechanism involves mediating the intracellular sequestration and active efflux of cisplatin, thereby reducing the effective drug dosage that reaches the nuclear DNA. The ability of CSGDA to independently identify this key efflux pump demonstrates that the framework can validate existing knowledge and has the potential to uncover implicit biological patterns, offering valuable leads for discovering novel resistance targets.

## 4. Conclusion

In this work, we propose CSGDA, a cell state-guided dynamic graph learning framework designed to resolve the generalization bottlenecks in single-cell drug response prediction. Our results suggest that in high-heterogeneity scenarios, such as cross-metastasis prediction, relying solely on static feature similarity often leads to unreliable drug sensitivity assessments. CSGDA addresses this by reconstructing graph topology through biological semantic guidance, which effectively reduces domain discrepancy and overcomes the noise sensitivity inherent in traditional KNN approaches. By prioritizing semantic consistency, this dynamic mechanism not only maintains high precision in baseline tasks but also significantly enhances AUPR and stability in complex cross-tissue transfer scenarios. Furthermore, our approach exhibits robust interpretability, accurately identifying key resistance drivers amidst distribution shifts. To more comprehensively address the complexities of the tumor microenvironment beyond transcriptomics, extending this framework toward multi-omics integration represents an essential future direction.

